# Intranasal administration of split-inactivated influenza virus vaccine mixed with a novel adjuvant elicits protective antibodies against seasonal influenza viruses

**DOI:** 10.1101/2025.10.23.684226

**Authors:** Michael A. Carlock, Katherine L. Cupo, Ted M. Ross

## Abstract

Seasonal influenza remains a persistent global health concern, and despite the availability of vaccines, there are both limitations in vaccine effectiveness and needle-associated hesitancy that reduce vaccine uptake. To address these challenges, this study evaluated the intranasal delivery of Fluzone, a commercially licensed split-inactivated influenza vaccine, formulated with or without adjuvant. Mice were vaccinated intranasally with Fluzone alone or combined with either Infectimune, a cationic lipid nanoparticle adjuvant, or TRAC478, a synthetic dual toll-like receptor liposome adjuvant containing TLR4 and TLR7/8 agonists. In immunologically naïve mice, a higher inoculation volume was more effective at eliciting protective antibody responses even though the same dose of vaccine antigen was used for vaccination. Fluzone mixed with adjuvant enhanced hemagglutination-inhibition (HAI) titers and all mice were protected against influenza virus challenge. Mice with pre-existing anti-influenza virus immunity that was elicited by prior infection with historical influenza strains had modest antibody responses and full protection following immunization with unadjuvanted vaccine. However, the inclusion of an adjuvant with Fluzone significantly raised HAI titers and HA-specific IgG levels. Pre-immune mice vaccinated with Fluzone mixed with either adjuvant had low to undetectable viral lung titers following challenge. Overall, intranasally delivered split-inactivated influenza vaccines paired with a mucosal adjuvant is a promising strategy to enhance vaccine effectiveness.

## INTRODUCTION

Influenza virus infection is a persistent public health challenge and causes serious respiratory illness each year, accounting for an estimated 290,000 – 650,000 global deaths [1, 2] and hospitalizations that strain healthcare systems and economies worldwide [3, 4]. Seasonal influenza vaccination is currently the most effective approach for preventing influenza-associated illnesses [5, 6]. Many people do not receive a seasonal influenza vaccine due to a variety of reasons, including affordability and access, vaccine hesitancy, perceptions that the vaccine is ineffective, or may actually cause influenza-like illness [7, 8]. Influenza viruses are unique due to their segmented, negative-sense RNA genome and an error-prone RNA polymerase that lacks proofreading capability [9-11]. This combination drives a high rate of point mutations (antigenic drift) and facilitates genetic reassortment (antigenic shift) when co-infection of diverse strains enables a “mix-and-match” of genomic segments, potentially leading to new, highly pathogenic variants that may result in global pandemics [12]. These evolutionary pressures are most pronounced in the major surface glycoproteins, hemagglutinin (HA) and neuraminidase (NA). Remarkably, these proteins can accommodate extensive amino acid substitutions without appreciable loss of viral fitness, thus allowing influenza viruses to evade existing antibody responses [13-15]. Current efforts are underway to improve upon seasonal influenza vaccines through new technologies [16, 17], including vaccines designed utilizing the computationally optimized broadly reactive antigen (COBRA) methodology employed by our lab [18-22]. However, other strategies to improve upon the current standard of care have also been implemented, such as high-dose [23] or adjuvanted [24] formulations for the elderly.

Fluzone (Sanofi Pasteur, Swiftwater, PA, USA), a split-inactivated virus vaccine that is administered to people via intramuscular injection, is among the most widely administered influenza vaccine each season [25, 26]. Fluzone is relatively inexpensive to produce compared to other influenza vaccine manufacturing platforms because it is generated from embryonated chicken eggs [27, 28]. Fluzone contains influenza virus particles that have been chemically inactivated using formaldehyde and chemically disrupted using a non-ionic surfactant, Triton^®^ X- 100 [29]. In addition to further concentration and purification steps, this process removes the internal viral RNA and proteins but leaves the HA and NA surface antigens to yield a vaccine product that is immunogenic and safe [30].

The goal of this study was to combine the safety of split-inactivated vaccines and the efficacy of adjuvanted formulations with the accessibility of intranasal vaccination to provide a protective vaccine formulation that could be administered intranasally. Mice were vaccinated with Fluzone alone or mixed with one of two novel adjuvants. Infectimune (R-DOTAP) is a cationic lipid nanoparticle (LNP) adjuvant that has been used in various animal studies [31-37]. By using positively charged LNPs to encapsulate negatively charged nucleic acids, the platform takes advantage of electrostatic attraction to enhance association with antigen-presenting cells and promote efficient uptake and intracellular delivery [33]. Pairing this adjuvant strategy with Fluzone should improve immune outcomes by delivering influenza antigens directly to immune cells in lymphoid tissues and stimulate more robust humoral and T-cell responses while minimizing inflammation.

The second novel adjuvant, TRAC478, is a synthetic dual toll-like receptor (TLR) liposome delivery system combining a TLR4 agonist and TLR7/8 agonist [38-40]. TLR4 is a pattern recognition receptor (PRR) that resides in the cell membrane and specifically recognizes lipopolysaccharide (LPS), the main cell wall component of Gram-negative bacteria [41, 42]. TLR4 agonists activate antigen-presenting cells by engaging the MyD88- and TRIF-dependent pathways that ultimately drive the production of pro-inflammatory cytokines for improved T-cell priming [43] as well as enhanced humoral responses [44]. TLR7 and TLR8 each primarily recognize ssRNA within the endosome [41]. TLR7 is highly expressed on plasmacytoid dendritic cells and B cells where it drives the production of IFN-α for antiviral immunity [41]. TLR8 is highly expressed on myeloid dendritic cells and monocytes where it stimulates pro-inflammatory cytokines such as IL□12 and TNF□α for the enhancement CD8+ T cell responses [41]. Combining TLR7 and TLR8 agonists enhances dendritic cell activation and antigen presentation, resulting in a robust Th1 immune response [41, 45, 46]. Co-encapsulating TLR4 and TLR7/8 within liposomes results in synergistic innate and adaptive immune responses characterized by a balanced Th1/Th2 response and longer-lasting protection [40, 47, 48]. Incorporating TRAC478 into Fluzone should engage innate signaling to drive more rapid and sustained humoral and cellular immune responses— resulting in substantially enhanced potency compared to unadjuvanted Fluzone.

Overall, this study demonstrated that intranasal mucosal delivery of a licensed human inactivated virus vaccine can enhance systemic IgG responses and elicit protective antibody titers. Vaccinated mice had anti-HA binding antibodies with HAI activity and survived influenza viral challenge following intranasal delivery of vaccine with or without adjuvant. Using an intranasal route of administration could broaden the use of this effective vaccine formulation for people.

## MATERIALS AND METHODS

### Influenza viruses

Influenza virus seed stocks were obtained through the Centers for Disease Control (CDC), the Influenza Reagents Resource (IRR), BEI Resources, or provided by Virapur (San Diego, CA, USA). Viruses were passaged once in the same growth conditions as they were received, either embryonated chicken eggs or semi-confluent Madin-Darby canine kidney (MDCK) cell cultures as per the instructions provided by the World Health Organization (WHO) [49]. Virus lots were HA confirmed with 0.8% turkey erythrocytes and made into aliquots for single-use applications.

The H1N1 viruses used in this study include A/Singapore/6/1986 (SG/86), A/Brisbane/02/2018 (BR/18), A/Victoria/2570/2019 (VC/19), A/Hawaii/70/2019 (HI/19), and A/Victoria/4897/2022 (VC/22). The H3N2 viruses used in this study include A/Panama/2007/1999 PN/99), A/Hong Kong/2671/2019 (HK/19), and A/Darwin/9/2021 (DR/21). The influenza B viruses used in this study include the Yamagata lineage B/Phuket/3073/2013 (B/PH/13) and the Victoria lineage B/Austria/1359417/2021 (B/AU/21).

### Mouse vaccination and infection

DBA/2J mice (females, 6–8 weeks old) were purchased from The Jackson Laboratory (Bar Harbor, ME, USA). Mice were housed in microisolator units and allowed free access to food and water. All animals were cared for under the USDA guidelines for laboratory animals, and all procedures were approved by the Cleveland Clinic Florida Research and Innovation Center Institutional Animal Care and Use Committee (IACUC) (protocol no. 2935). All animal work is additionally performed in accordance with the National Research Council (NRC) Guide for the Care and Use of Laboratory Animals, the Animal Welfare Act, the CDC/NIH Biosafety in Microbiological and Biomedical Laboratories, and in accordance with the ARRIVE guidelines.

Immunologically naïve mice were randomly divided into six groups (n = 15/group) where they were intranasally immunized with commercially available 2022-2023 Fluzone (Sanofi Pasteur, Swiftwater, PA, USA) split inactivated influenza virus vaccine (**Fig. 1A**). Mice were administered ∼1μg/HA (12μL) or ∼0.5μg/HA (6μL) of Fluzone, or ∼1μg/HA in a 24μL dose with sterile phosphate-buffered saline (PBS; Corning, Tewkbury, MA, USA), or ∼1μg/HA in a 24μL dose with a cationic lipid nanoparticle adjuvant known as R-DOTAP or Infectimune (PDS Biotechnology Corporation, Florham Park, NJ, USA), or ∼0.5μg/HA in a 24μL dose with Infectimune and sterile PBS (**Table 1**). A group receiving 24μL of sterile PBS only was additionally included for a mock control. For the adjuvanted groups, ∼72μg of Infectimune was used (12μL from a 6mg/mL stock). Vaccines were administered at day 0 and again at day 21 (**Fig 1A**). Blood was collected from the facial vein at 14- and 23-days post-boost. Sera was isolated from the blood via centrifugation at 6000 rpm for 5 min in BD Microtainer blood collection tubes (BD, Franklin Lakes, NJ, USA). Clarified serum was combined for the 14- and 23-day timepoints. 5 days after the final bleed (28 days post-boost), each vaccination group was divided into three infection groups (n=5/group) and challenged with HI/19 H1N1 at 1×10^5 PFU/50μL, or 1×10^4 PFU/50μL, or 1×10^e3 PFU/50μL diluted in sterile PBS. Following infection, weights for each mouse were recorded daily, and all animals were monitored for 11 days for clinical signs which included labored breathing, lethargy, failure to respond to stimuli, severe respiratory distress, or weight loss, with 25% used as the primary determination of humane endpoint.

**FIG 1.**
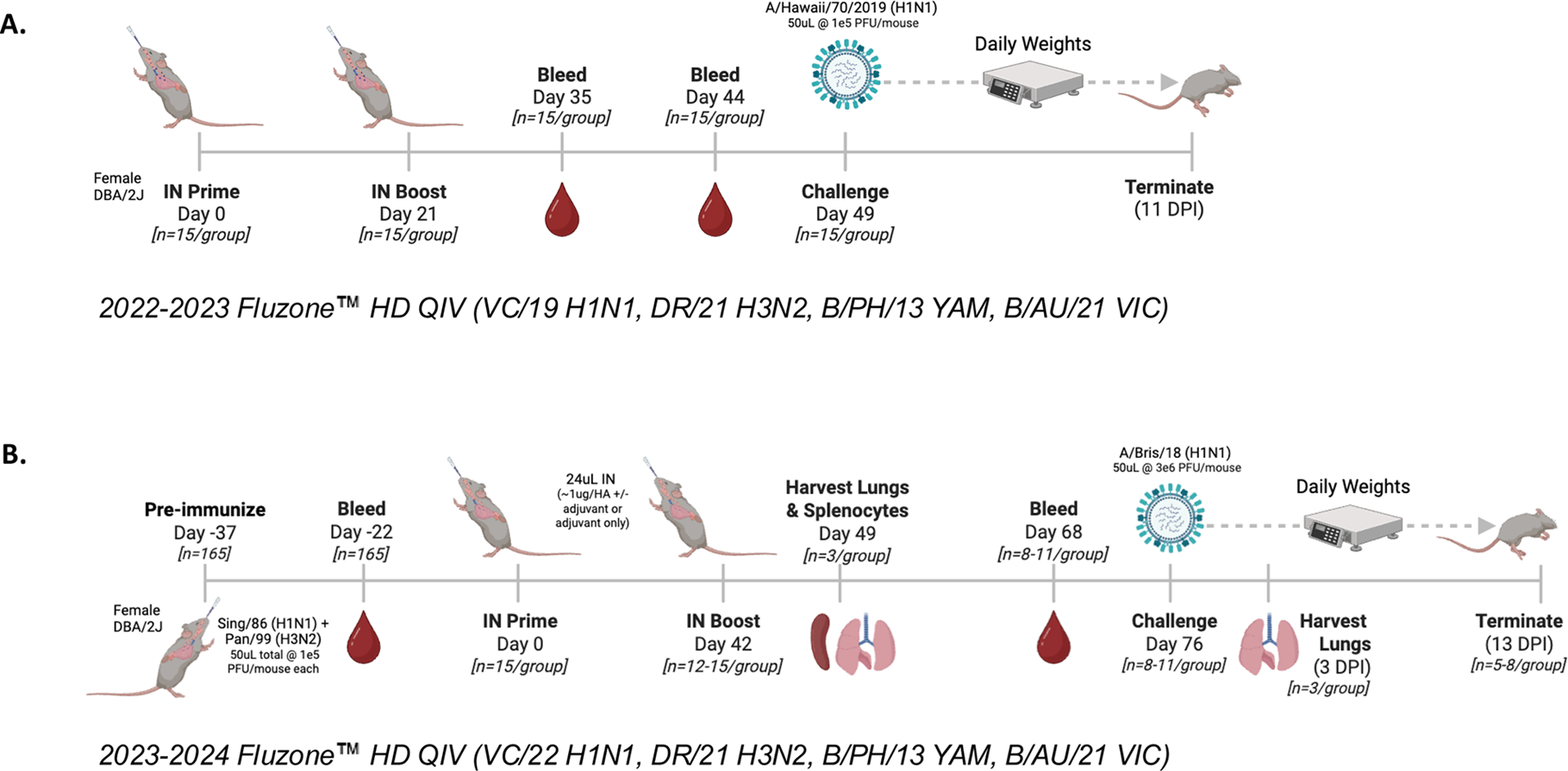
Study regimen. Influenza-naïve mice (A) were intranasally vaccinated (n=15/group) at days 0 and 21, bled at days 35 and 44, and then challenged at day 49 with HI/19 H1N1 virus at 1e5 PFU/50µL. Pre-immune mice (B) were infected with historical H1N1 (SG/86) and H3N2 (PN/99) viruses at day -37, bled at day -22 to confirm seroconversion, and then intranasally vaccinated (n=12-15/group) at days 0 and 42. Lungs and splenocytes were collected at day 49, blood was collected at day 68, and then mice were challenged at day 76 with BR/18 H1N1 virus at 3e6 PFU/50µL. Lungs were collected 3 days post-infection. Daily weights and clinical signs were recorded following either infection.

**TABLE 1.**
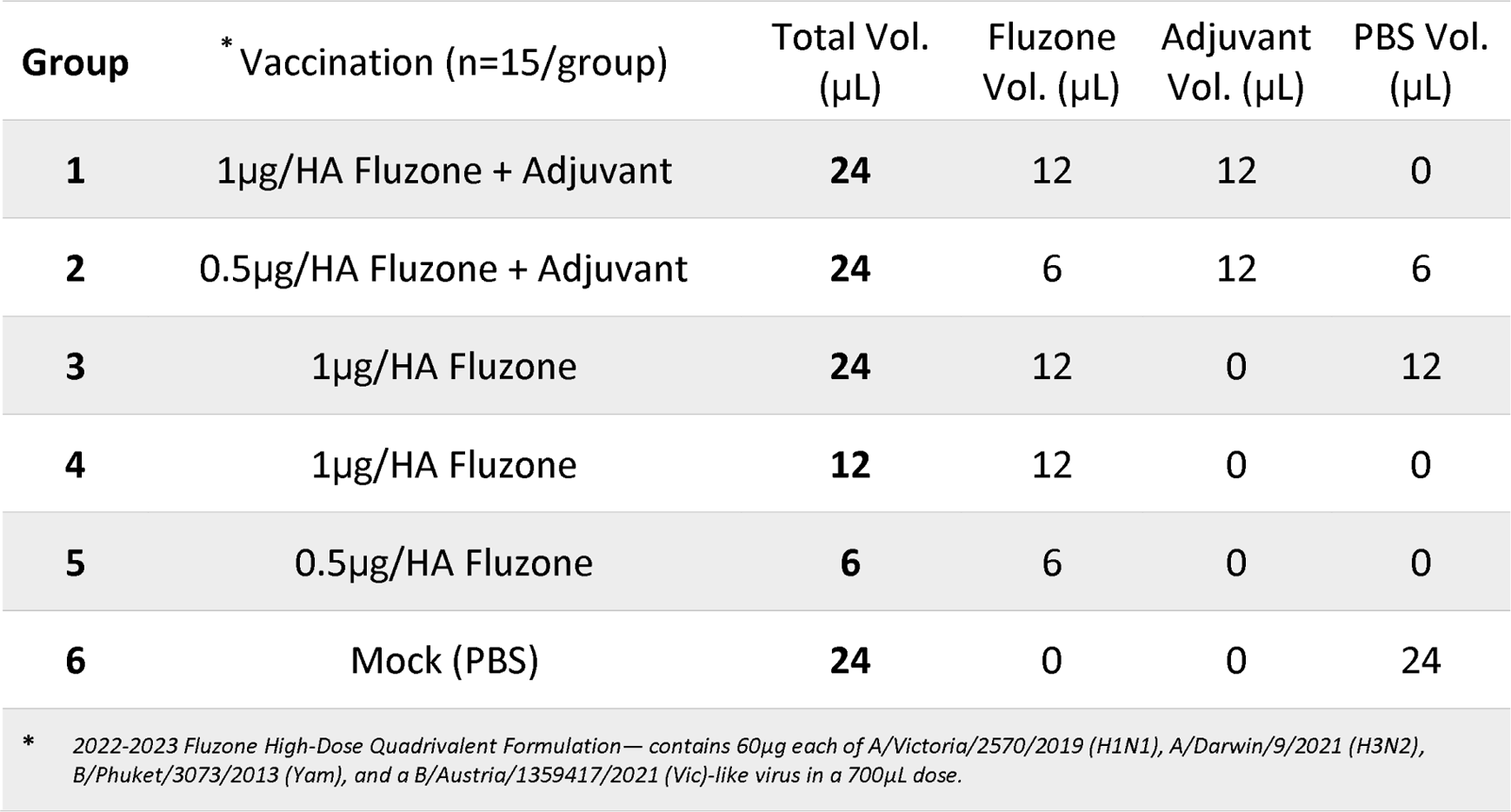
Vaccination Groups.

| Group | * Vaccination (n=15/group) | Total Vol. (μL) | Fluzone Vol. (μL) | Adjuvant Vol. (μL) | PBS Vol. (μL) |
| --- | --- | --- | --- | --- | --- |
| 1 | 1μg/HA Fluzone + Adjuvant | 24 | 12 | 12 | 0 |
| 2 | 0.5μg/HA Fluzone + Adjuvant | 24 | 6 | 12 | 6 |
| 3 | 1μg/HA Fluzone | 24 | 12 | 0 | 12 |
| 4 | 1μg/HA Fluzone | 12 | 12 | 0 | 0 |
| 5 | 0.5μg/HA Fluzone | 6 | 6 | 0 | 0 |
| 6 | Mock (PBS) | 24 | 0 | 0 | 24 |

\* 2022-2023 Fluzone High-Dose Quadrivalent Formulation — contains 60μg each of A/Victoria/2570/2019 (H1N1), A/Darwin/9/2021 (H3N2), B/Phuket/3073/2013 (Yam), and a B/Austria/1359417/2021 (Vic)-like virus in a 700μL dose.

**TABLE 1** Vaccination Groups
| **Group** | **\* Vaccination (n=15/group)** | **Total Vol.**<br>**(uL)** | **Fluzone**<br>**Vol. (uL)** | **Adjuvant**<br>**Vol. (uL)** | **PBS Vol.**<br>**(uL)** |
| --- | --- | --- | --- | --- | --- |
| **1** | 1ug/HA Fluzone + Adjuvant | **24** | 12 | 12 | 0 |
| **2** | 0.5ug/HA Fluzone + Adjuvant | **24** | 6 | 12 | 6 |
| **3** | 1ug/HA Fluzone | **24** | 12 | 0 | 12 |
| **4** | 1ug/HA Fluzone | **12** | 12 | 0 | 0 |
| **5** | 0.5ug/HA Fluzone | **6** | 6 | 0 | 0 |
| **6** | Mock (PBS) | **24** | 0 | 0 | 24 |
| **\* 2022-2023 Fluzone High-Dose Quadrivalent Formulation— contains 60ug each of A/Victoria/2570/2019 (H1N1), A/Darwin/9/2021 (H3N2), B/Phuket/3073/2013 (Yam), and a B/Austria/1359417/2021 (Vic)-like virus in a 700uL dose.** |  |  |  |  |  |

Additional mice were pre-immunized to influenza A virus using a mixture containing H1N1 (SG/86) and H3N2 (PN/99) at 1×10^5 PFU/50μL each, diluted in sterile PBS (**Fig. 1B**). Three weeks later, blood was collected from the facial vein and sera was isolated as previously described. Seroconversion to these two strains was confirmed for each mouse via hemagglutination inhibition (HAI) assay. Mice were then randomly divided into five groups (n = 15/group) where they were intranasally immunized with commercially available Fluzone (2023-2024 high-dose, quadrivalent formulation) in a 24μL dose diluted in half with sterile PBS (approximately 1μg/HA), or in a 24μL dose diluted in half with one of two adjuvants: Infectimune, or TRAC478 (Inimmune, Missoula, MT, USA), a synthetic dual TLR liposome delivery system combining a TLR4 adjuvant (INI-2002) and a TLR7/8 adjuvant (INI-4001). Mice were vaccinated with Fluzone (1μg based upon HA content) with or without adjuvant. Mock vaccinated mice were administered a 24μL dose divided in half with sterile PBS and either adjuvant. Mice were vaccinated at day 0 and boosted at day 42. Each vaccination consisted of TRAC478 containing ∼1.13μg INI-2002 and 2.4μg INI-4001 in a 6μL volume plus and 6μL of sterile PBS (18μL PBS for adjuvant-only). At day 49 days (7 days post-boost) spleens and lungs (n=3/group) were collected in B cell media (1L RPMI supplemented with 100mL FBS, 20mL HEPES, 20mL of a 50x MEMAA solution, 10mL of a 100x NEAA solution, 10mL Penicillin-Streptomycin, 10mL sodium pyruvate, 2g of sodium bicarbonate, and 3.5μL of 2-mercaptoethanol). Splenocytes were processed and stored in liquid nitrogen and lungs were homogenized and stored at -80°C. At 4 weeks post-boost, blood was collected and sera isolated as described above. At day 76, mice were challenged with BR/18 H1N1 influenza virus (3×10^6 PFU/mouse/50μL). At day 79 post-infection, lungs were collected (n=3/group) to assess viral lung titers. Lungs were collected on dry ice and then stored at -80°C. Following infection, mice were assessed for morbidity and weight loss, as well as mortality for 14 days.

### Hemagglutination-inhibition (HAI) assay

The HAI assay was used to assess functional antibodies to the HA that prevent the agglutination of avian red blood cells (RBCs) by H1N1, H3N2, and influenza B viruses (IBVs). The protocols used for this assay were adapted from the World Health Organization (WHO) laboratory influenza virological surveillance manual [49]. Sera samples were treated with receptor destroying enzyme (RDE) (Denka, Seiken, Co., Japan) to inactivate nonspecific inhibitors, according to the manufacturer’s instructions prior to their use in the assay. In brief, three volumes of RDE were added to one volume of sera and incubated overnight at 37°C. The next day, the RDE was inactivated by incubating the samples at 56°C in a water bath for 30-60 min, after which 6 volumes of PBS were added to each sample, resulting in a final serum dilution of 1:10. RDE treated sera was then diluted in a series of two-fold serial dilutions in 96-well V-bottom plates, and an equal volume of influenza virus, adjusted to 8 hemagglutination units (HAU)/50μL diluted in PBS, was added to each well of the plate. The plates were allowed to incubate at room temperature for 20 min. After incubation, 50μL of a solution consisting of 0.8% turkey RBCs (Lampire Biologicals, Pipersville, PA, USA) diluted in PBS was added to each well. The plates were then mixed by gentle agitation and allowed to incubate for another 30 min at RT. After incubation with RBCs the plates were tilted to observe the hemagglutination inhibition. The HAI antibody titer was determined by taking the reciprocal dilution of the last well that contained non-agglutinated RBCs. Positive control ferret reference serum was also included to confirm assay consistency between runs. Prior to use, the turkey RBCs were washed twice with 1xPBS, stored at 4°C, and used within 24 h of preparation. IBVs used in HAI assays were first treated with diethyl ether, as previously reported [50].

### Anti-HA Enzyme Linked Immunosorbent Assay (ELISA)

ELISAs were used to independently assess the elicited serum Ig subclass IgG1and IgG2a antibody titers against a panel of influenza H1, H3, and influenza B HA antigens. In brief, each well of an Immulon 4HBX 96-well flat-bottom plate (Thermo Fisher, Waltham, MA, USA) was coated with 100mL of carbonate buffer (pH 9.4) containing 5mg/mL of bovine serum albumin (BSA) (Thermo Fisher, Waltham, MA, USA), and 500ng/mL of full-length recombinant HA protein that was transiently expressed and purified as previously described [51]. Recombinant HA proteins used to coat plates for this assay included: WI/19 H1 (EPI_ISL_404460), VC/22 H1 (EPI_ISL_17830834), DR/21 H3 (EPI_ISL_2233238), B/PH/13 (Yamagata lineage) (EPI_ISL_165882), and B/AU/21 (Victoria lineage) (EPI_ISL_1519459). Plates were then incubated overnight at 4°C in a humidified chamber. The next day, the plates were decanted and blocked with 200mL/well blocking buffer (1xPBS containing 5% BSA, 2% bovine gelatin, and 0.05% Tween-20 at 37°C for 90 minutes in a humidified chamber. Serum samples were serially diluted 3-fold in blocking buffer starting at a dilution of 1:100. Plates were then incubated overnight at 4°C in a humidified chamber. The next day, the plates were washed 5 times with wash buffer (1xPBS, 0.05% Tween-20). The wash buffer was then decanted, and secondary antibody diluted 1:4000 in blocking buffer was added to each well. Goat Anti-Mouse IgG1 (cat #1070-05) and Goat Anti-Mouse IgG2a (cat #1083-05) HRP-conjugated antibodies from Southern Biotech (Birmingham, AL, USA) were used. The plates were incubated at 37°C for 90 minutes in a humidified chamber and then washed 5 times with wash buffer. The plates were then decanted and 100μL/well of ABTS Diammonium Salt (Amersco, Solon, Ohio, USA) working solution was added to each plate. Plates were incubated at 37°C for 15 minutes, and then 50μL/well of a 1% sodium dodecyl sulfate solution was added to each well to stop the colorimetric reaction. Optical density values of each well were measured at 414nm using a spectrophotometer (PowerWave XS, Agilent Technologies, Santa Clara, CA, USA), and plotted in GraphPad Prism (San Diego, CA, USA) for each dilution point. Total area-under-the-curve (AUC) for each vaccination point was obtained, as were endpoint dilution titers.

### Viral Plaque Assay

MDCK cells were seeded at a concentration of 1×10^6 cells/well into six-well plates on day one. The following day, after MDCK cells reached ∼90% confluency in each well, the plates were washed three times with Dulbecco’s modified Eagle medium supplemented with 1% penicillin-streptomycin (DMEM + P/S). Nasal wash samples were thawed on ice and serially diluted 10-fold in DMEM + P/S. Media was removed from the plates and 100μL of the viral dilutions were added. The plates incubated at room temperature for 1 h, but gently rocked every 15 min. Following the incubation, the plates were washed twice with DMEM + P/S. Following the second wash, an overlay solution of plaque media (2x minimum essential medium, HEPES, L-glutamine, P/S, sodium bicarbonate, and 1 μg/mL of TPCK-treated trypsin) mixed 1:1 with 1.6% agarose was added to each well. Plates were then incubated at 37°C + 5% CO_2_ for 72 h. Afterwards, the gel overlays were removed, and the cells were fixed with 10% buffered formalin for 10 min and then stained with 1% crystal violet for 10 min, all at room temperature. Plates were then rinsed thoroughly with fresh water to remove excess crystal violet, and plates were allowed to air dry for at least 24 h. Plaques were then counted, and the viral titer (PFU per milliliter) for each nasal wash sample was calculated using the number of plaque colonies and the dilution factor.

### Statistical Methods

Statistical significance was defined as a p value < 0.05. GraphPad Prism version 10 software was used throughout. Bar graphs show geometric mean titer ± geometric standard deviation and were analyzed using an ordinary one-way ANOVA with Tukey’s multiple comparisons test to compare the mean of each column with the mean of every other column. For ELISA curves and weight loss curves, a two-way ANOVA mixed-effects model with the Geisser-Greenhouse correction and Tukey’s multiple comparisons test was utilized.

## RESULTS

### Intranasal administration of split-inactivated influenza virus vaccine elicits protective antibodies against seasonal influenza viruses

The commercial split-inactivated influenza virus vaccine Fluzone (Sanofi Pasteur, Swiftwater, PA, USA) is approved for intramuscular administration. In this study, mice were administered Fluzone (2022-2023 QIV) intranasally, with or without an adjuvant, to determine the effectiveness of delivering a split-inactivated influenza virus vaccine to the respiratory mucosa (**Table 1**). Against the H1N1 vaccine virus, VC/19, mice vaccinated with Fluzone plus the Infectimune adjuvant had, on average, hemagglutinin-inhibition (HAI) activity ≥1:40 regardless of the dose of vaccine administered (**Fig. 2A**). In contrast, mice vaccinated without adjuvant had, on average, HAI titers <1:40, with significantly lower titers for the group administered only 6μL of Fluzone (**Fig. 2A**). There was low HAI activity detected in the sera against the H3N2 vaccine component, DR/21 (**Fig. 2B**). In contrast to the HAI activity against the influenza A virus (IAV) components, Fluzone vaccinated mice had, on average, HAI titers ≥1:40 against the influenza B virus (IBV) components regardless of the inclusion of adjuvant (**Fig. 2C-D**). Mice vaccinated with adjuvanted Fluzone had on average 2-4 fold higher HAI titers against IBV than mice vaccinated without adjuvant; however, this serum HAI activity was statistically higher than mice administered vaccine in a 6μL or 12μL volume (**Fig. 2C-D**).

**FIG 2.**
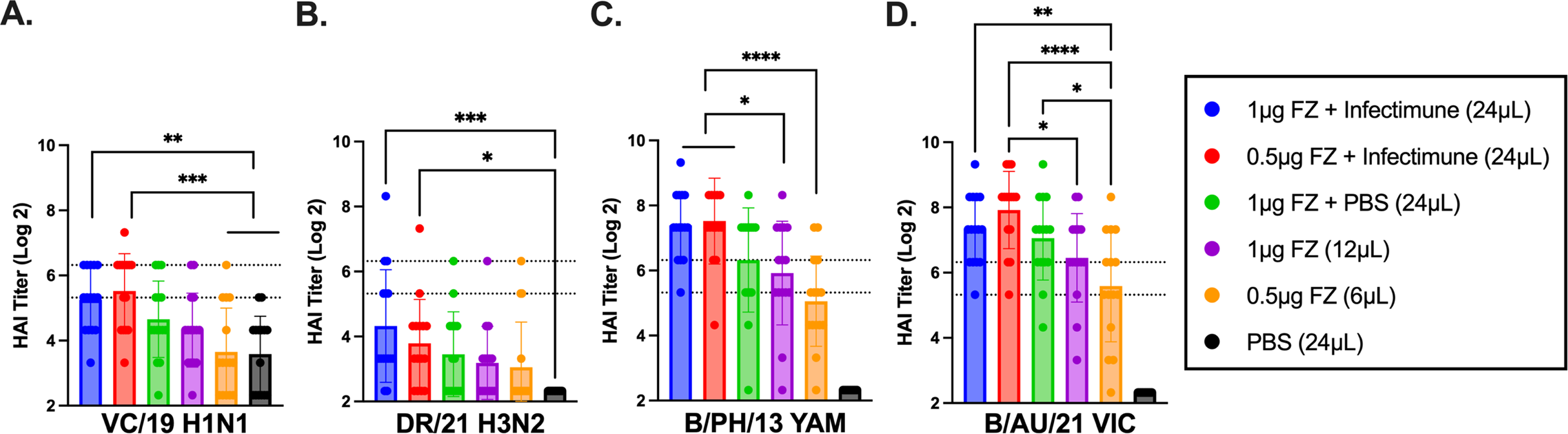
HAI serum antibody titers induced by intranasal vaccination with Fluzone at different volumes and antigen concentration. HAI titers were assessed against the vaccine strains included in the 2022-2023 Fluzone vaccine following. Values are the log2 HAI titers of each individual mice (n=15/group) from antisera collected following two intranasal vaccinations. Dotted lines indicate a 1:40 to 1:80 HAI titer range. Statistical analyses were performed using one-way ANOVA to compare the means of each column: p > 0.05 = ns, p ≤ 0.05 = *, p ≤ 0.01 = **, p ≤ 0.001 = ***, p ≤ 0.0001 = **** (note: stats for the PBS group against IBVs in panels C and D not shown for easier readability)

Post-vaccination serum samples were additionally evaluated for anti-HA IgG1 and IgG2a isotypes using enzyme linked immunosorbent assay (ELISA) (**Fig. 3**). Serum from mice immunized with adjuvanted Fluzone had consistently the highest responses, particularly for the group administered a 1μg/HA dose of vaccine that elicited relatively high levels against DR/21 whereas the other vaccinated groups did not (**Fig. 3C**). For the unadjuvanted groups, 1μg/HA elicited similar serum responses no matter if mice received 12μL or 24μL (**Fig. 3**). Mice receiving a 6μL dose had much lower responses, but they also only received 0.5μg/HA. And there was no detectable serum antibodies from mice immunized with only PBS. In general, low levels of IgG2a were observed for any of the vaccinated groups against any of the vaccine strains. However, the adjuvanted groups had slightly higher levels, led by the group immunized with 0.5μg/HA Fluzone; as such, this group had slightly lower ratios of IgG1 to IgG2a.

**FIG 3.**
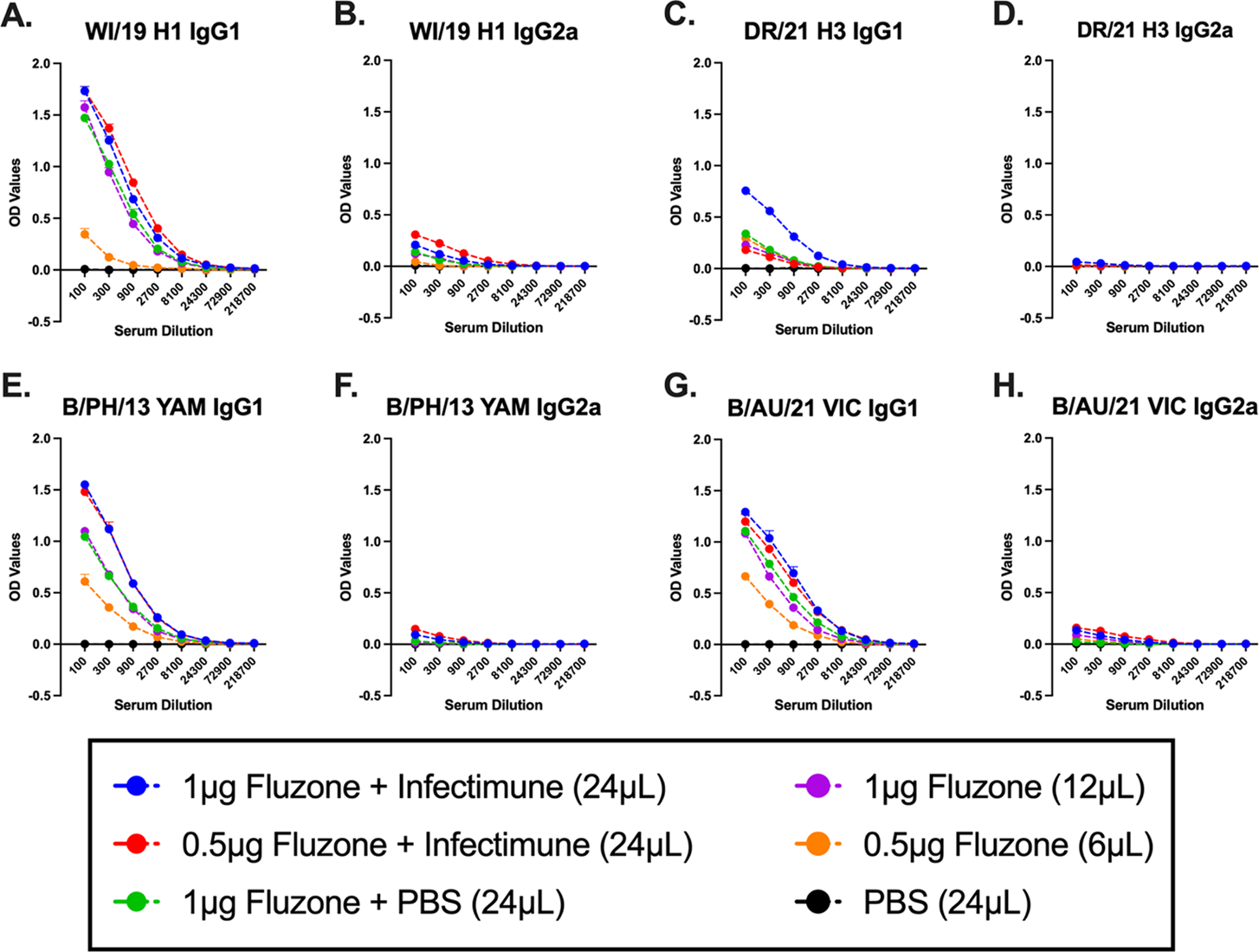
ELISA serum antibody titers induced by intranasal vaccination with Fluzone at different volumes and antigen concentration. ELISA OD values shown at each serum dilution for IgG1 (A, C, E, G) and IgG2a (B, D, F, H) in response to WI/19 H1 (A-B), DR/21 H3 (C-D), B/PH/13 Yam (E-F), and B/AU/21 Vic (G-H).

At day 49, mice were challenged with the H1N1 virus, HI/19 (**Fig. 4**). Following infection, mock-vaccinated mice rapidly lost body weight, and by day 6 post-infection reached humane endpoint after losing ≥25% original body weight (**Fig. 4A**) with no mice surviving challenge (**Fig. 4B**). Mice vaccinated with Fluzone without adjuvant had similar weight loss as mock vaccinated mice; however, mice vaccinated with 1μg of Fluzone had 20-80% survival depending on the inoculation volume (**Fig. 4B**). In contrast, all adjuvanted mice survived infection. Mice vaccinated with 1μg of Fluzone plus Infectimune lost an average of 7% of their original body weight by day 2 post-infection and slowly began to regain weight over the next 10 days (**Fig. 4A**). Mice vaccinated with 0.5μg of Fluzone plus Infectimune lost an average of 15% of original body weight by day 4 post-infection and then maintained that weight for the next 10 days of observation (**Fig. 4A**).

**FIG 4.**
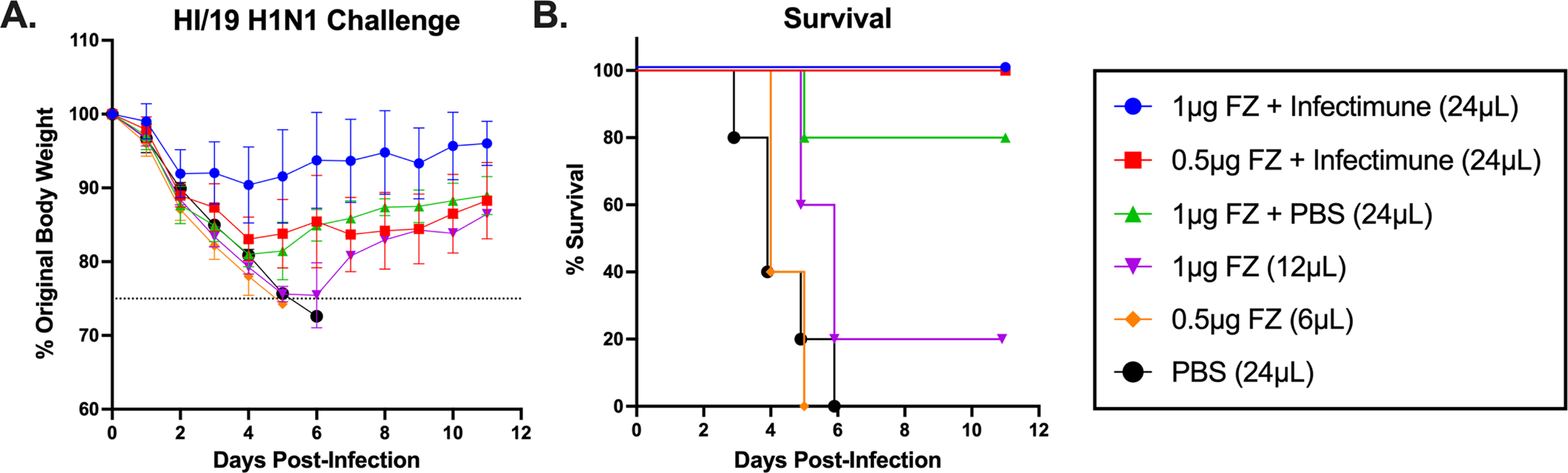
Weights and survival following challenge. Four weeks following the final vaccination, mice were infected with A/Hawaii/70/2019 H1N1 at 1×10^5 PFU/mouse. Percent original weight of mice (A) and survival (B) is shown over an 11-day period.

Collectively, these initial findings support the potential of intranasal Fluzone as a promising mucosal vaccination strategy—especially when formulated with an adjuvant. In mice, this strategy triggered strong humoral responses and significantly improved survival following H1N1 challenge. Notably, inoculation volume played a decisive role: mice receiving 1µg Fluzone in a larger volume (24µL vs. 12µL) exhibited greater protection, despite similar HAI titers— suggesting that delivery parameters may influence mucosal efficacy, perhaps by enhancing antigen distribution or retention in the nasal passages.

### A synthetic dual TLR liposome adjuvant increases protective antibodies elicited by intranasal Fluzone vaccination

To further determine the effectiveness of Fluzone vaccine administered intranasally, another set of mice was pre-immunized to historical influenza A viruses (**Fig. 1B**). Briefly, mice were infected with a mixture of SG/86 (H1N1) and PN/99 (H3N2) and then allowed to rest for one month prior to vaccination. Mice were subsequently vaccinated twice with Fluzone (2023-2024 QIV) formulated with or without adjuvant. Adjuvanted groups received one of two adjuvants: Infectimune, as described in the previous section, or TRAC478, a synthetic dual TLR liposome delivery system combining TLR4 and TLR7/8 agonists.

Following the second vaccination, pre-immune mice vaccinated with unadjuvanted Fluzone had an average HAI titer ≥1:40 against the two H1N1 viruses tested, BR/18 and VC/22 (**Fig. 5A-B**). These same sera had lower HAI activity against the two H3N2 viruses tested, with little to no HAI activity against HK/19 (**Fig. 5C**) and an average HAI titer just at 1:40 against DR/21 (**Fig. 5D**). Fluzone vaccinated mice had higher HAI titers against the influenza B viruses with an average HAI titer ≥1:160 (**Fig. 5E-F**). Pre-immune mice immunized with Fluzone plus either Infectimune (FZL+LInfectimune) or TRAC478 (FZL+LTRAC478) adjuvants exhibited higher serum HAI titers on average compared to mice administered unadjuvanted Fluzone (Fluzone + PBS) (**Fig. 5**). Notably, mice vaccinated with Fluzone plusLTRAC478 had statistically higher serum HAI titers against the H1N1 strains compared to mice vaccinated with unadjuvanted Fluzone or vaccination with Fluzone plus Infectimune (**Fig. 5A–B**). In general, mice vaccinated with Fluzone plus adjuvant had slightly elevated HAI titers against the H3N2 strains (**Fig. 5C–D**). Although mice vaccinated with Fluzone plus TRAC478 had statistically higher titers against HK/19 with the average HAI titers elicited in mice vaccinated with unadjuvanted Fluzone or Fluzone plus Infectimune had HAI titers less than 1:40 (**Fig. 5C**). Lastly, mice vaccinated with Fluzone plusLTRAC478 had significantly higher serum HAI titers against B/PH/13 compared mice vaccinated with Fluzone or Fluzone plus Infectimune and higher HAI titers against B/AU/21 compared to mice vaccinated with unadjuvanted Fluzone (**Fig. 5E–F**).

**FIG 5.**
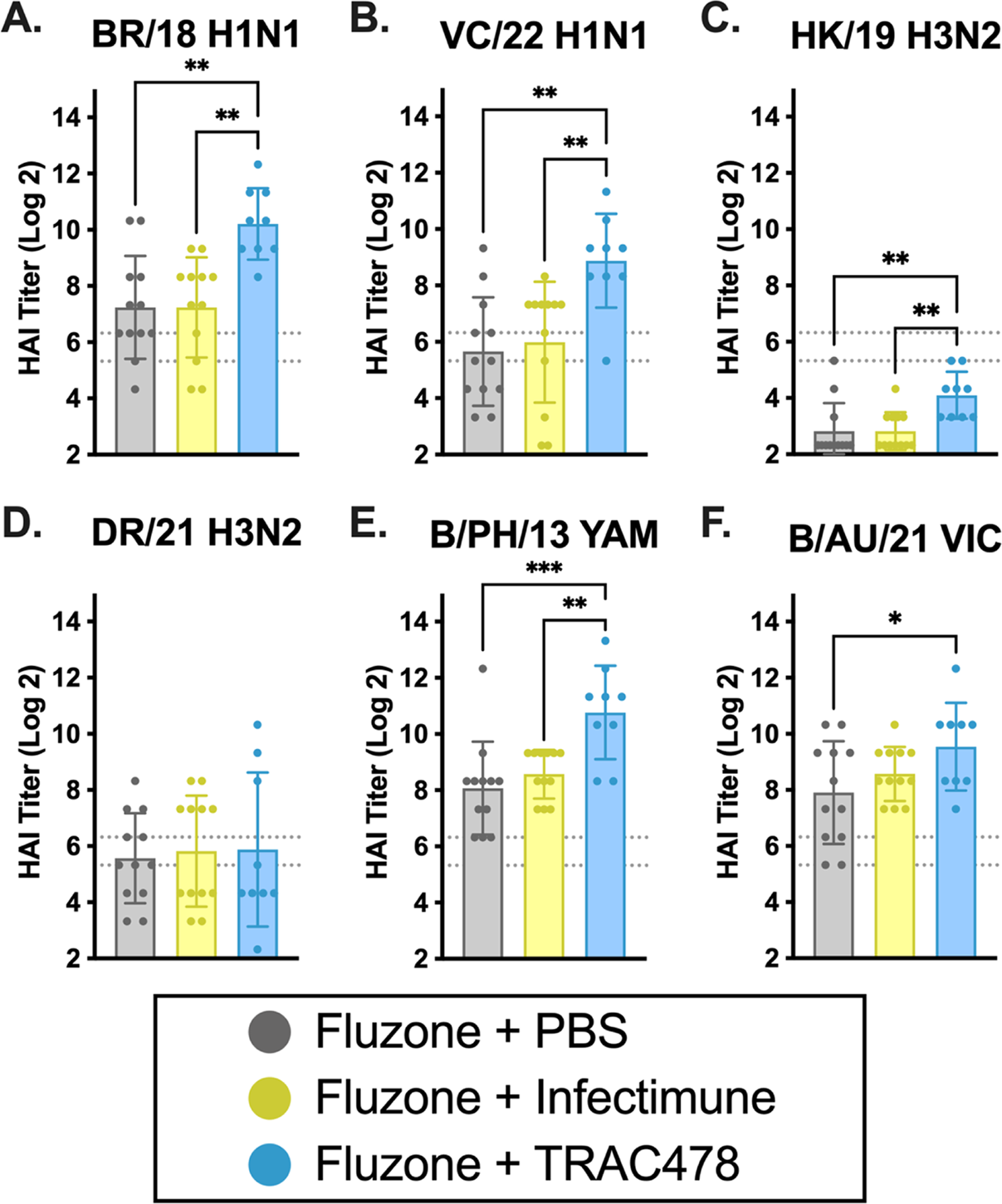
HAI serum antibody titers induced by adjuvanted and non-adjuvanted Fluzone seasonal influenza vaccination. HAI titers were assessed against two H1N1 viruses (A-B), two H3N2 viruses (C-D), one Yamagata virus (E), and one Victoria virus (F). Adjuvant-only mice (not shown) were all HAI negative against each virus. Log2 HAI titers for individual mice are shown as scatter dot plots with bars representing the geometric mean titer of each group and error bars indicating the geometric standard deviation. Dotted lines indicate a 1:40 to 1:80 HAI titer range. Statistical analyses were performed using one-way ANOVA to compare the means of each column: p > 0.05 = **ns**, p ≤ 0.05 = **\***, p ≤ 0.01 = **\*\***, p ≤ 0.001 = **\*\*\***, p ≤ 0.0001 = **\*\*\*\***

In general, mice vaccinated with unadjuvanted Fluzone or Fluzone plus Infectimune had higher serum IgG1 against the H1 and influenza B HA proteins than mice vaccinated with Fluzone plus TRAC478 (**Fig. 6**). These same mice had weak anti-HA titers against the H3 HA protein DR/21, regardless of if the vaccine was mixed with adjuvant (**Fig. 6C-D**). Overall, mice had lower serum IgG2a titers against any of the HA proteins compared to IgG1 titers, except against the H3 HA. In general, mice vaccinated with Fluzone mixed with TRAC478 had higher IgG2a serum anti-HA titers compared to unadjuvanted Fluzone resulting in a more mixed T helper response (**Fig. 6I**). In contrast, mice vaccinated with Fluzone plus Infectimune had more robust IgG1 titers compared to IgG2a titers with a more predominate T helper 2 response.

**FIG 6.**
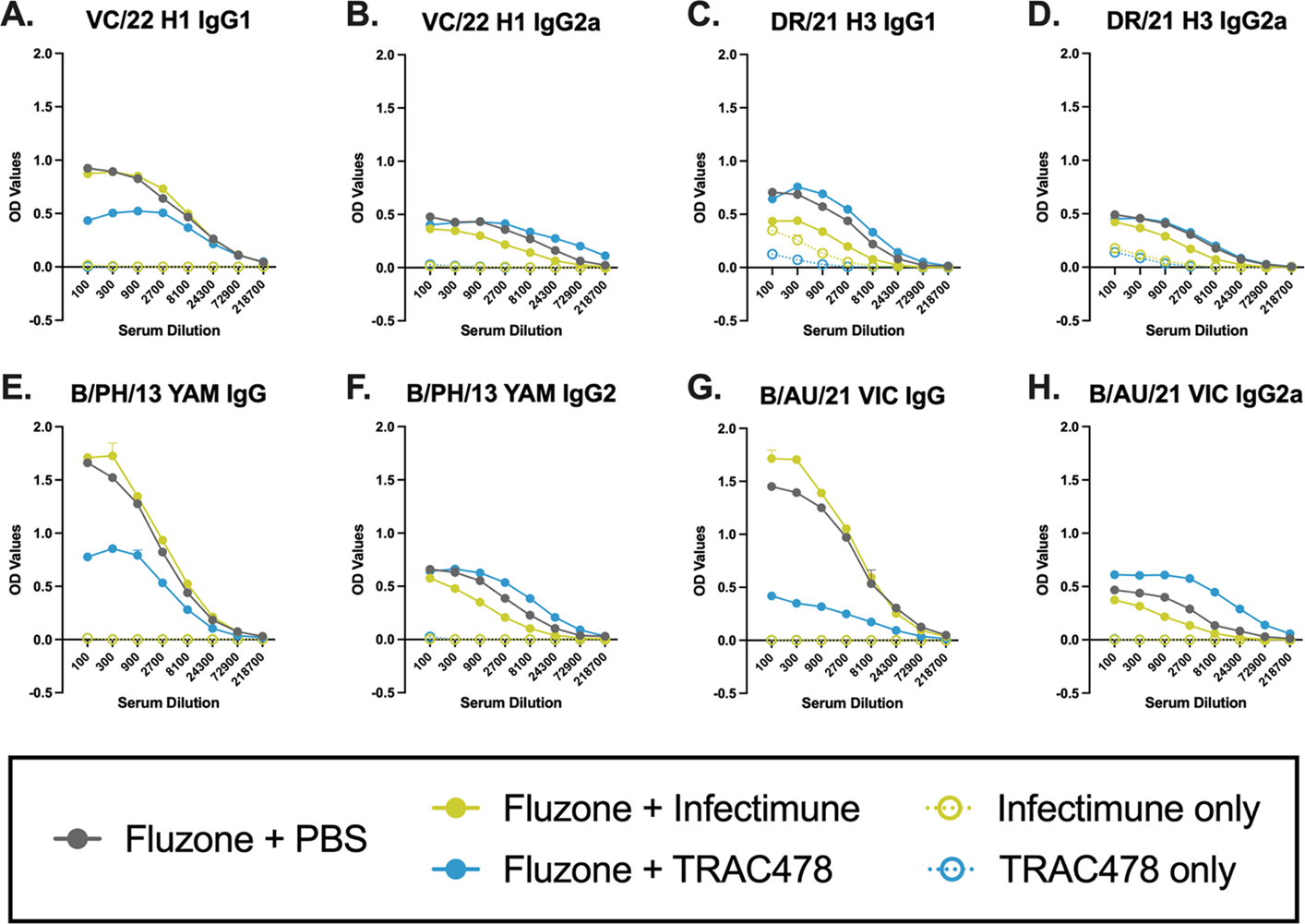
ELISA serum antibody titers induced by adjuvanted and non-adjuvanted Fluzone seasonal influenza vaccination. ELISA OD values shown at each serum dilution for IgG1 (A, C, E, G) and IgG2a (B, D, F, H) in response to WI/19 H1 (A-B), DR/21 H3 (C-D), B/PH/13 Yam (E-F), and B/AU/21 Vic (G-H).

Mice were challenged with the BR/18 (H1N1) influenza virus 34 days following the final vaccination (**Fig. 7**). Mice vaccinated with only adjuvant rapidly lost weight and, on average, had 80% of their original body weight by day 5 post-infection (**Fig. 7A**) with 60-70% of the mice surviving challenge (**Fig. 7B)**. Then, these mice slowly regained weight over the 14 days of observation. In contrast, all vaccinated mice administered Fluzone with or without adjuvant survived challenge with little weight loss. On average, vaccinated mice lost ∼5-7% weight (**Fig. 7A**) that was statistically higher between days 2-7 compared to mice that were administered only adjuvant (**Suppl Table 1**). At day 3 post-challenge, lungs were harvested (n=4/group) and assessed for viral lung titers (**Fig. 7C**). Mice vaccinated with only adjuvant had statistically higher viral titers (∼5×10^5 pfu/mL) than mice vaccinated with Fluzone with or without adjuvant. No virus was detected in any mouse, except for one animal that was vaccinated with Fluzone plus Infectimune (**Fig. 7C**).

**FIG 7.**
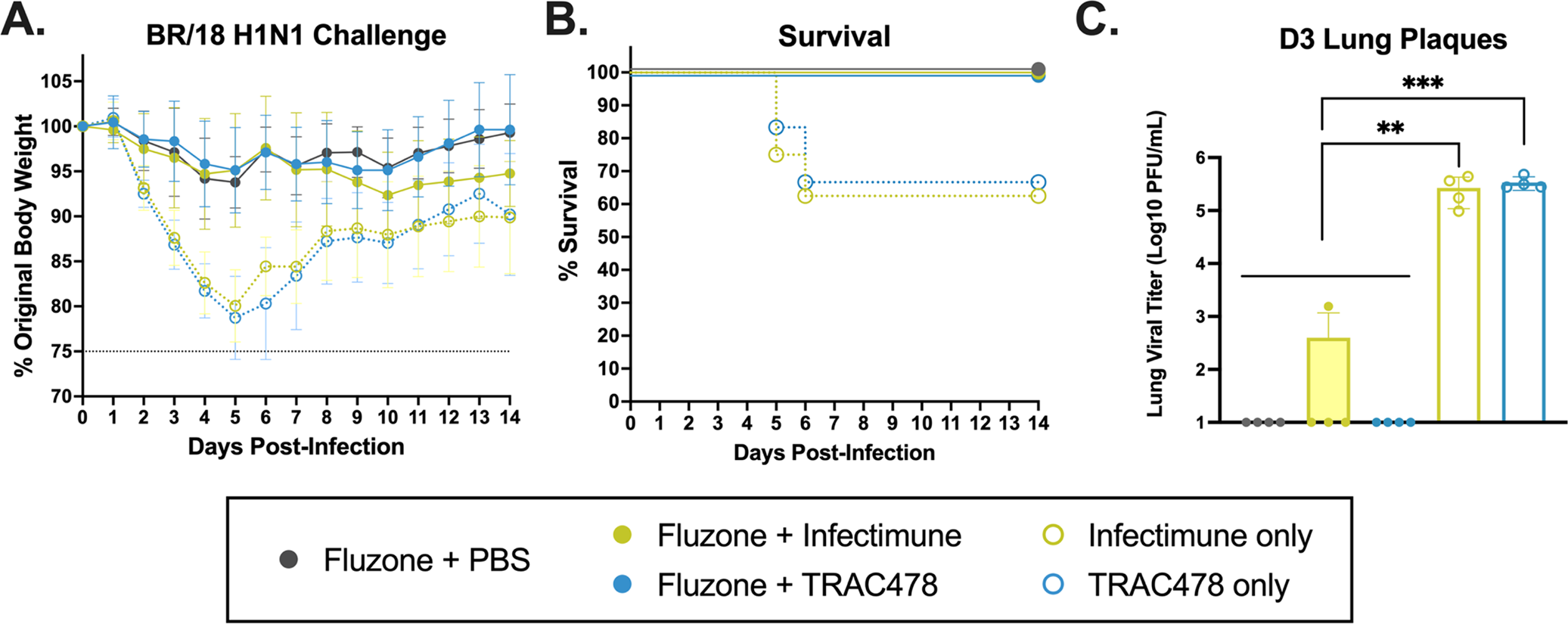
Weights, survival, and lung plaques following challenge. Challenges are performed 27 days post-vaccination with A/Brisbane/02/2018 H1N1 virus @ 3e6 PFU/mouse. Mice are monitored for weights and clinical symptoms daily, and lungs are collected 3 days post-infection. Percent original weight of mice (A) and survival (B) is shown over a 14-day period. Viral plaques on lungs (C) collected from 3 mice from each group is shown with statistical analyses performed using one-way ANOVA to compare the means of each column: p > 0.05 = ns, p ≤ 0.05 = *, p ≤ 0.01 = **, p ≤ 0.001 = ***, p ≤ 0.0001 = ****

## DISCUSSION

Intranasal administration of influenza vaccines offers the dual advantage of needle-free delivery and the potential to stimulate both mucosal and systemic immune responses, which are critical for protection against respiratory pathogens. Recently, the U.S. Food and Drug Administration (FDA) approved an at-home intranasal administration of the live-attenuated influenza vaccine FluMist [52]. In this study, the split-inactivated quadrivalent Fluzone vaccine that is approved for intramuscular administration was used intranasally vaccinate mice. Intranasal delivery of Fluzone elicited detectable serum HAI activity, particularly administered with an adjuvant. In immunologically naïve mice, the inclusion of Infectimune, a cationic liposomal adjuvant, boosted HAI titers and increased survival following challenge compared to unadjuvanted Fluzone. In mice with pre-existing immunity to influenza A viruses, the immune responses elicited by unadjuvanted Fluzone and Fluzone mixed with Infectimune were statistically similar. In contrast, Fluzone formulated with TRAC478 induced the strongest HAI titers and balanced anti-HA IgG1 to IgG2a responses. These findings suggest that Infectimune promotes a T helper 2-skewed response while TRAC478 appears to support a more balanced T helper 1/T helper 2 profile.

Pre-immune mice vaccinated with Fluzone with or without adjuvant were protected from an H1N1 influenza virus challenge. In naïve mice, adjuvanted Fluzone provided complete protection from H1N1 influenza virus challenge. Intranasal Fluzone without adjuvant was less effective at eliciting protective immune responses, but only if the volume administered was >6μL. In mice, intranasal administration of a fluid volume less than 10μL is typically used to limit antigen delivery to the upper respiratory tract of mice [53, 54]. This suggests that the increased volumes used in this study would reach the lower respiratory tract. The lower respiratory tract is more susceptible to immune-mediated tissue damage; therefore, vaccines that reach the lower respiratory tract should be examined carefully. In contrast, initial immunizations with the TRAC478 adjuvant resulted in 17% of mice dying following vaccination. However, mice boosted with the same dose and volume (24μL) of vaccine, but half the amount of adjuvant, all survived vaccination with no observable vaccine-mediated complications. While TRAC478 was highly immunogenic, it appears to be more toxic when used at larger amounts but can be formulated for safety.

Translating this approach to humans would pose even greater formulation and dosing challenges. Intramuscular vaccines, such as Fluzone, are not optimized for nasal retention or stability since vaccine antigens need to survive the nasal mucus, which can dilute or trap vaccine antigens, as well as nasal enzymes can degrade proteins [55]. Relatively large doses or specialized delivery systems (*e.g.* mucoadhesive carriers, nasal sprayers, or potent adjuvants) are often needed for efficient uptake, and only a few human-safe mucosal adjuvants exist [56, 57]. Safety and regulatory issues are also nontrivial. For example, Nasalflu (Berna Biotech, Bern, Switzerland), which is an intranasal influenza vaccine that uses a virosome carrier system and is adjuvanted with an *E. coli* heat labile toxin, was withdrawn from the market after an association with Bell’s palsy was observed [58, 59].

In this study, Fluzone administered intranasally effectively elicited protective immune responses in mice with and without the use of an adjuvant. Each season, Fluzone elicits neutralizing antibodies in younger people using a standard dose of antigen, but it is less effective in elderly populations. To increase effectiveness in older adults, two strategies have been employed to enhance immune responses. One approach is to increase the dose of the vaccine from 15μg to 60μg per antigen (Fluzone High Dose) [23] and the second approach has been the addition of an adjuvant (FluAd produced by Seqirus, Holly Springs, NC, USA) [24]. Both approaches enhance the effectiveness of split-inactivated influenza vaccines in older populations. Neither of these vaccines are administered intranasally to people, however, clinical studies could repurpose standard dose split-inactivated virus vaccines mix with adjuvants. These influenza vaccines administered via a different delivery route, such as intranasally, could more effectively elicit protective immune responses, particularly in high-risk populations. As presented in this report, the addition of an adjuvant that has greater potential to stimulate adverse events following vaccination may not be warranted via intranasal delivery. Of course, any intranasal version of a licensed split-inactivated influenza virus vaccine would require full clinical testing and regulatory review as a new product. Still, this study highlights the potential of using split-inactivated influenza virus vaccines via intranasal delivery.

## Data Availability

Data is publicly available via Import database.

## Acknowledgements

Some of the influenza viruses were obtained through the International Reagent Resource (IRR), Influenza Division, WHO Collaborating Center for Surveillance, Epidemiology, and Control of Influenza, Centers for Disease Control and Prevention (Atlanta, Georgia, USA). We also would like to thank the University of Georgia and Cleveland Clinic Animal Resource staff, technicians, and veterinarians for their excellent animal care.

## Author Contributions

M.A.C., K.L.C., and T.M.R. conceptualized the experiments. M.A.C. and K.L.C. conducted the animal work and collected samples. M.A.C. and K.L.C. performed assays. M.A.C. prepared the figures and analyzed the data. M.A.C. and T.M.R. wrote and edited the manuscript. All authors read and approved the final version of the manuscript.

---

**SUPPL TABLE 1.**
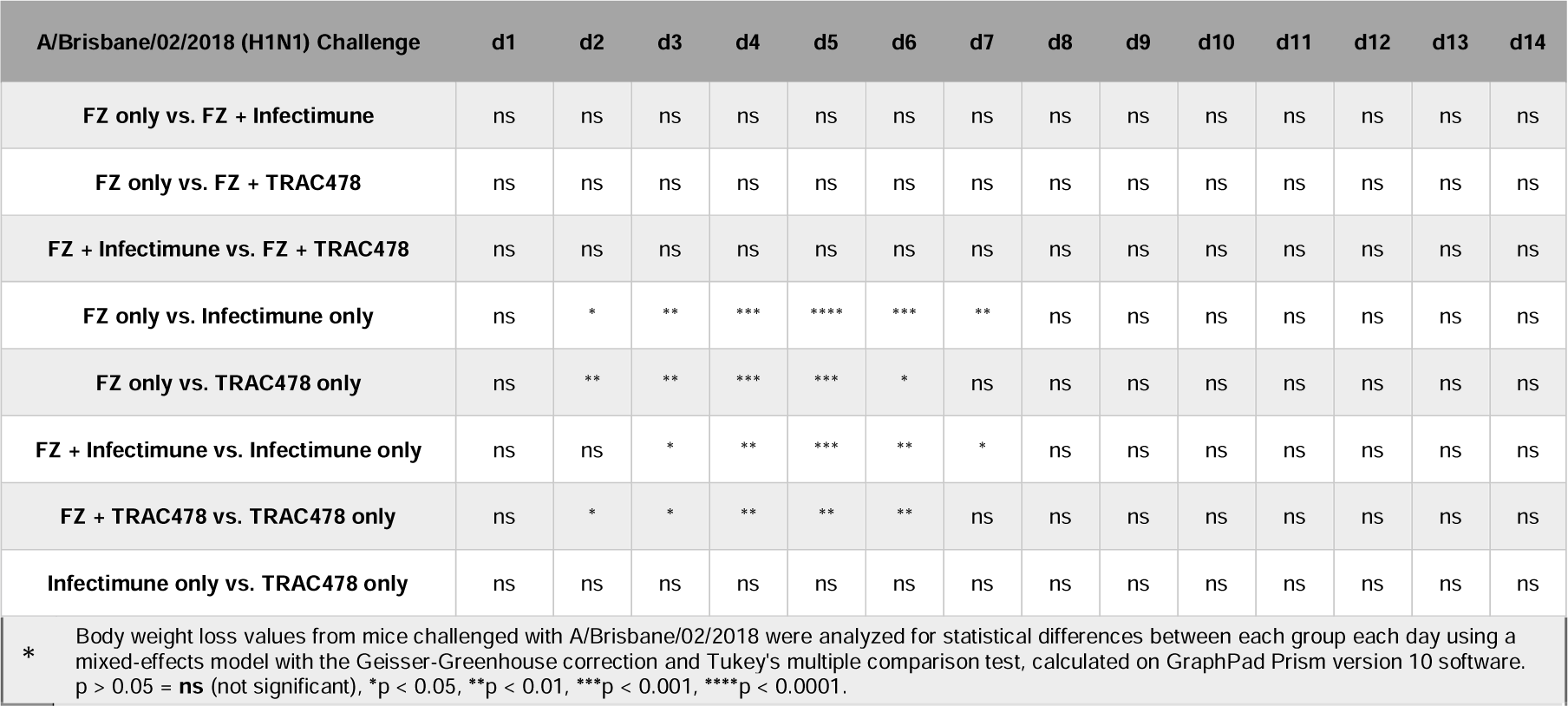
Statistical difference of body weight loss between mouse vaccine groups*.

