## Supplementary material for "Intranasal administration of split-inactivated influenza virus vaccine mixed with a novel adjuvant elicits protective antibodies against seasonal influenza viruses": Supplmental Table 1

**SUPPL TABLE 1** Statistical difference of body weight loss between mouse vaccine groups*

| A/Brisbane/02/2018 (H1N1) Challenge | d1 | d2 | d3 | d4 | d5 | d6 | d7 | d8 | d9 | d10 | d11 | d12 | d13 | d14 |
|---|---|---|---|---|---|---|---|---|---|---|---|---|---|---|
| **FZ only vs. FZ + Infectimune** | ns | ns | ns | ns | ns | ns | ns | ns | ns | ns | ns | ns | ns | ns |
| **FZ only vs. FZ + TRAC478** | ns | ns | ns | ns | ns | ns | ns | ns | ns | ns | ns | ns | ns | ns |
| **FZ + Infectimune vs. FZ + TRAC478** | ns | ns | ns | ns | ns | ns | ns | ns | ns | ns | ns | ns | ns | ns |
| **FZ only vs. Infectimune only** | ns | * | ** | *** | **** | *** | ** | ns | ns | ns | ns | ns | ns | ns |
| **FZ only vs. TRAC478 only** | ns | ** | ** | *** | *** | * | ns | ns | ns | ns | ns | ns | ns | ns |
| **FZ + Infectimune vs. Infectimune only** | ns | ns | * | ** | *** | ** | * | ns | ns | ns | ns | ns | ns | ns |
| **FZ + TRAC478 vs. TRAC478 only** | ns | * | * | ** | ** | ** | ns | ns | ns | ns | ns | ns | ns | ns |
| **Infectimune only vs. TRAC478 only** | ns | ns | ns | ns | ns | ns | ns | ns | ns | ns | ns | ns | ns | ns |

* Body weight loss values from mice challenged with A/Brisbane/02/2018 were analyzed for statistical differences between each group each day using a mixed-effects model with the Geisser-Greenhouse correction and Tukey's multiple comparison test, calculated on GraphPad Prism version 10 software. $p > 0.05$ = **ns** (not significant), *$p < 0.05$, **$p < 0.01$, ***$p < 0.001$, ****$p < 0.0001$.
